# Homology of the dark cells of Paleozoic liverworts with the specialized oil body cells of modern liverworts (Marchantiophyta)

**DOI:** 10.64898/2025.12.24.696392

**Authors:** Susan Tremblay, Josep Mercadal

**Affiliations:** Department of Integrative Biology, University of California, Berkeley, California, 94720, USA; Department of Plant Developmental Biology, Max Planck Institute for Plant Breeding Research (MPIPZ), 50829 Cologne, Germany

**Keywords:** Marchantiophyta, *Metzgeriothallus*, plant fossils, liverwort oil bodies, organelles

## Abstract

- Oil bodies are a synapomorphy of liverworts (Marchantiophyta), a major group of land plants with a sparse fossil record. Paleozoic liverworts sometimes possess dark cells that appear similar in distribution to liverwort oil body cells. The Middle Devonian *Metzgeriothallus sharonae* provides an opportunity for comparison with modern liverworts.
- Shale samples were collected and the carbonaceous fossils isolated by acid maceration. Museum shale specimens of *Pallaviciniites devonicus* were also obtained and processed. The relative location, composition, frequency and spatial distribution of the fossil dark cells was compared to that of oil cells in extant taxa.
- Chemical characterization of the fossil dark cells is consistent with cells that contained lipids, the main component of oil bodies. Quantitative analyses revealed that the frequency and spatial distribution of dark cells are comparable to oil body cells. The dark cells of *M. sharonae* show clumping near the thallus edge, suggesting an anti-herbivore function.
- These results support the hypothesis that the dark cells of Paleozoic liverworts and the oil body cells of extant lineages are homologous structures with a shared developmental origin, providing a new character that can be used to assign fossils and that sheds light on the evolution and function of liverwort oil bodies.

## Introduction

Liverworts (Marchantiophyta) are a morphologically and ecologically diverse group of land plants. Early cladistic studies generally resolved liverworts as sister to the remaining embryophytes (Mishler & Churchill, 1984; Kenrick & Crane, 1997; Qui *et al*. 2006). More recent large molecular data sets have supported a sister group relationship between mosses and liverworts (Cox *et al.,* 2014; Wickett *et al.,* 2014; Puttick *et al.,* 2018; Bechteler *et al*., 2023). While the earliest (Silurian) land plant body fossils have been assigned to the vascular plants, much of the earlier Ordovician evidence of embryophytes—mainly enigmatic spores and cellular fragments—is thought to have liverwort affinities (Gensel *et al*., 1990; Edwards *et al*., 1995; Strother, 2000; Graham *et al*., 2004), and the earliest mesofossils with bryophytic affinities have been placed in the liverwort lineage (Hueber, 1961; Lacey, 1969; Krassilov & Schuster, 1984; Oostendorp, 1987; Hernick *et al*., 2008). All six major lineages of extant liverworts are thought to have diverged by the end of the Paleozoic (Heinrichs et al. 2007; Bechteler *et al*., 2023). Unfortunately, until recently, the only Paleozoic mesofossils that have been placed unequivocally in the liverwort clade were represented by a small number of specimens that exhibited only vegetative characters, which are, as in all embryophytes, prone to homoplasy (Krassilov & Schuster, 1984).

Because of their sparse early fossil record (Tomescu *et al.,* 2018), early diversification and character evolution in liverworts remains poorly understood (Wiens 2004; Crandall-Stotler *et al*., 2005). One roadblock has been assigning fossils to liverworts with so few characters available. Until now, one of the most important synapomorphies of the group, their unique organelles called oil bodies, has never been used to assign fossils. The ability to do so would be especially advantageous as oil bodies vary in a number of characters that are taxonomically informative within the liverwort clade (Schuster, 1966; Crandall-Stotler *et al*., 2005). Understanding of cellular patterns in early liverworts could also facilitate interpretation of the cellular scraps that pre-date the first land plant mesofossils, the so-called ’waifs and strays’ (Wellman, 2003). This could provide information about occurrence of land plants and timing of appearance of characters (Edwards *et al*., 1995).

Oil bodies are true, membrane-bound organelles. Present in 90% of liverwort species across all main lineages, they have been secondarily lost in several groups (Schuster, 1966; He *et al*., 2013). They sequester lipophilic compounds (Asakawa, 1995), and Suire (2000) has shown evidence of terpenoid synthesis in the oil bodies of *Marchantia polymorpha*. Varying in size, texture, and color, they often impart characteristic odors to a particular species. The ultrastructure, chemical composition, and taxonomic significance of liverwort oil bodies have been extensively investigated (Schuster, 1966; Asakawa, 2004) yet their exact function was unknown until recently. A growing body of research has found evidence of anti-microbial properties of oil bodies (Millar *et al.,* 2004) and has increasingly supported an anti-herbivory function (He *et al*., 2013; Romani, 2020). In the vast majority of taxa oil bodies are found in all cells of the plant and depending on the taxon, an individual liverwort cell may contain one to as many as fifty or more oil bodies. In the Marchantiopsida and the Treubiales (Haplomitriopsida) they are only found in specialized cells–sometimes referred to as idioblasts–that are nearly filled with a single large oil body, while other cells are oil body-free.

The first Paleozoic fossil liverworts were described by Walton, who named five taxa from the Middle and Upper Carboniferous (Walton,1925; Walton, 1928). The first Devonian liverwort, *Pallaviciniites devonicus*, was described by Hueber (1961) and was accepted as the oldest fossil liverwort, until the spectacularly well-preserved *Metzgeriothallus sharonae* was discovered occurring in large numbers in a lens of dark gray shales in a Middle Devonian outcrop in New York State (Hernick *et al*., 2008). All these Devonian and Carboniferous taxa were discovered in fine shales and mudstones and despite their relative rarity are usually very well-preserved, exhibiting fine cellular detail (Walton, 1925 & 1928; Hueber, 1961; Hernick *et al*., 2008).

A notable feature of many of these fossils is the presence of dark, scattered cells that appear denser and more opaque than the surrounding cells (Fig. 1A-D). These cells have been described as possible storage cells (Hernick *et al*., 2008) or the site of waste accumulation or secretory cells (Walton, 1925). Schuster (1966) compared them to the oil body cells found in some extant lineages, an explanation that seems most likely if a homologous structure in modern groups is to be proposed. Other types of specialized cells/structures can occur in liverworts, such as endogenous gemmae (asexual propagules) in the Fossombroniacaeae but these are found only on occasion in some individual plants and are confined to specific parts of the plant (Crandall-Stotler pers. comm.). As far as specialized cells that are ubiquitous and occur on all parts of the plant, the only example in extant liverworts are the specialized oil body cells found in the Treubiales and the Marchantiales (Fig. 1E-H). Although Schuster placed the Carboniferous taxon *Treubiites kidstonii* in the modern Treubiaceae based on other characters, he did note that the presence of scattered dark cells supported the placement (Schuster, 1984). Initial observations of the *M. sharonae* fossils suggest that the dark cells are more numerous near the edge of the unistratose thallus wings, a situation also occurring in at least some cases of modern liverworts; for example, in asexual propagules of *Marchantia polymorpha* oil body cells appear to be more frequent around the perimeter of the disc-like structures (Labandeira *et al*., 2013; Romani, 2020, Fig. 5F).

**Figure 1.**
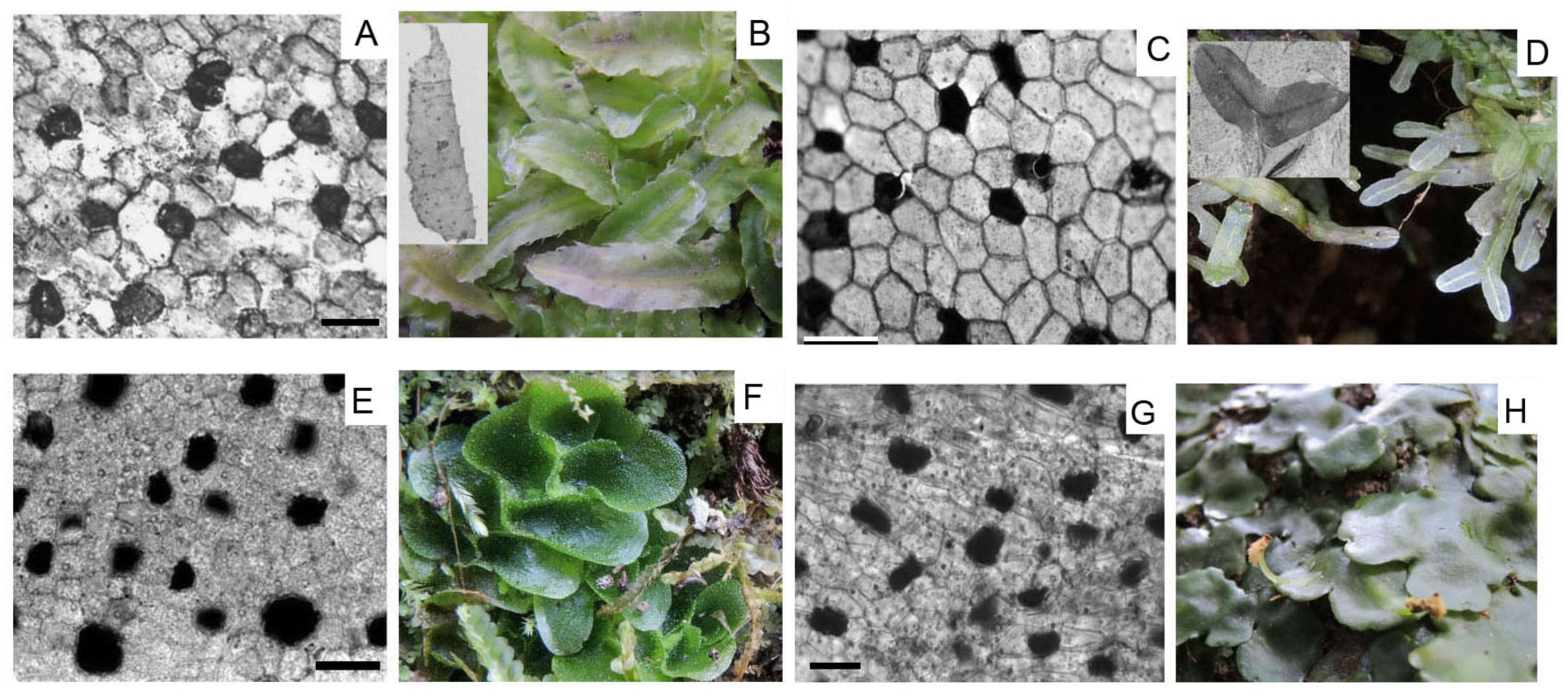
Extant oil body cells and fossil ’dark cells’ with respective habit photographs of taxa used in the quantitative analyses. Top row: Cells and thallus of fossil taxa *Pallaviciniites devonicus* (Hueber) Schuster, (A and inset of B) and *Metzgeriothallus sharonae* (C and inset of D) with representatives of the modern families they have been assigned to: *Pallavicinia xiphoides* (B) and *Metzgeria furcata* (L.) Dumort. (D). Bottom row: Cells and habit photos of extant liverworts *Treubia lacunosa* (Colenso) Prosk. (E, F, UC 2036933) and *Monoclea forsteri* Hook. (G, H, UC 2091243). Because cell size tends to vary among liverworts, the four cell photos (A,C,E,G) have been slightly adjusted for scale to show a similar number of cells. Scale=0.1mm.

Confirmation of this observation would lend support to the proposed function of oil bodies as anti-herbivory, as the edge of a foliar structure would be more likely to be attacked by predators (Labandeira *et al*., 2013).

By definition, homologous structures should be in the same relative position; be made of the same materials; and develop in the same way (Owen, 1848; Roth, 1988). In the case of fossils we can’t follow development very precisely, but the large numbers of fossil specimens of *M. sharonae* available provide an opportunity to compare the position and frequency of the cells to those of some modern taxa; and the remarkable preservation of the Cairo fossils allows us to take a closer look at the structure and composition of the dark cells themselves. As *M. sharonae* is the oldest generally accepted liverwort, occurring at a time when the main extant lineages may not all have yet diverged (Heinrichs et al. 2008) it could provide insight into the evolution of oil bodies and into the function of these unique organelles.

## Methods

### Sample collection and processing

Fossils: *Metzgeriothallus sharonae* material was collected from the Cairo Quarry, New York State, in August 2011 and June 2012. This locality is an old quarry located just south of New York State Route 145 at the Cairo Highway Department headquarters, 3.22 km NW of Cairo, Greene County, New York (42.32° N and 74.04° W, NAD 83). The material was collected from three locations approximately 3m apart along the 25m wide exposed lens of dark gray shales and mudstones where the liverworts were discovered. Unmacerated material (still in the shale matrix) of *Pallaviciniites devonicus* collected from the type locality by Fran Hueber was obtained from NMNH (Accession nr 348094) for the purposes of this study. Shale fragments were immersed in 49% hydrofluoric acid in order to dissolve the shale matrix. The isolated carbonaceous fossils were rinsed in water until a neutral pH was obtained, and then rinsed two more times. Thousands of carbonaceous *M. sharonae* fossils, mostly small fragments, were isolated. Selected specimens were dehydrated to a 50% glycerin solution, then the fossil fragments were mounted on glass slides and fixed with permanent slide mounting medium (Glycerin Gelatin). Over six hundred permanent slides were made. Specimens were photographed through a Nikon D90 digital camera mounted on a Leica DMRB compound microscope. High-resolution images of the mounted specimens were taken with a Leica DM2500 microscope using Differential Interference Contrast, a Plan Apo 63x Oil objective, and a Nikon DS-Fi1 Digital Camera. Extended depth of field images were generated using Adobe Photoshop’s stacking feature.

Fossil specimen selection for quantitative analyses: One hundred and twenty *Metzgeriothallus sharonae* samples were selected for the quantitative analyses based on the following criteria: unistratose thallus fragments larger than 1mm x 1mm that showed good preservation and included a portion of the midrib. Specimens were chosen from a variety of rocks from the three different locations along the layer. The selected specimens were permanently mounted on glass slides and imaged at 100 magnification and saved as TIF files, all at the same orientation, with the midrib partially visible along the lower edge. Closer examination of the photographs showed that many specimens had other dark types of structures in their cells, and many had variably fading dark cells in multiple shades of gray. Also, many specimen images included damaged areas. The original 120 photographs were eventually narrowed down to those specimens that had little damage, clearly defined dark cells, and little evidence of other cellular structures that could be confused with possible oil bodies, leaving 55 specimen photographs to use for quantitative analyses.

Extant taxa: *Targionia hypophylla* L. was collected in Sonoma County, California and *Marchantia polymorpha* gemmae were collected from the teaching collection housed at the UC Berkeley University greenhouse. Fresh material of *Treubia lacunosa* (Colenso) Prosk. was collected from four localities in New Zealand’s South Island and *Monoclea forsteri* Hook. from one Stewart Island locality. They were imaged while the material was still fresh at 100x magnification using a Leica compound microscope fitted with a digital camera. Voucher specimens of *T. lacunosa* and *M. forsteri* have been deposited in the University Herbarium, UC Berkeley, (specimen numbers UC2036933, UC2036941, UC2036942, UC2036943, UC2036944, UC2036945, UC2036946, and UC2091252). A small number of individual plants was collected from several areas at each locality. For the quantitative studies 2 - 6 undamaged ’leaf’ tips per *T. lacunosa* shoot were used, from the ’middle’ leaves only from each individual plant, mainly the unistratose region. Hand sections of *M. forsteri* were made along the plane of the upper and lower epidermis of the multistratose thalloid plants where most of the oil body cells occur using a single edged razor blade. The sections were hand cut as thinly as possible, generally 2-3 cells thick.

### SEM and stereo microscopy

Specimens still on the matrix were initially examined with a Leica Wild MZ8 stereo microscope using polarized fiber optic light source and a rotatable analyzer in order to increase contrast (Hernick *et al*., 2008). Water or cedarwood oil was applied to rock surfaces with a fine brush before photographing the compressions to help resolve details (Hernick *et al*., 2008). For higher magnifications a Nikon D90 digital camera mounted on a Leica DMRB compound microscope was used. Scanning electron microscopy (SEM) was done with the Hitachi TM-1000 SEM on uncoated specimens.

### Fluorescence microscopy

Mounted specimens were photographed with a Leica DM2000 epifluorescence microscope equipped with violet and UV excitation filter sets and a phototube, which fits a Nikon DS-Fi1 Digital Camera.

### Cell counts, morphology, and spatial distribution for fossil and extant taxa

We analyzed the number, morphology and spatial distribution of dark cells in *M. sharonae* and *P. devonicus*, and oil body cells in *T. lacunosa.* We first segmented a subset of the specimens (those where cell walls could be clearly distinguished) with the deep learning-based cell segmentation software cellpose (Stringer *et al*., 2021; Fig. 4A-C). We subsequently performed manual corrections of badly segmented cells. After segmentation, we annotated and labeled cell types (dark vs. light for *M. sharonae* and *P. devonicus*, and oil vs. non-oil body cell for *T. lacunosa*). Finally, we extracted positional (cell locations and cell neighborhoods) and morphological properties (cell area, perimeter, and cell wall lengths) from the labeled image. Because only three specimens of *Pallaviciniites devonicus* were well-preserved enough for analysis, all three were used and all cells (excluding the midrib) of each specimen were counted. The same pipeline was applied to the photographs of the ’leaf’ tips of *T. lacunosa* in order to use a similar area for the extant and extinct taxa. However, this meant that portions of the ’leaf’ that were more than one cell thick were included. Care was taken when imaging and counting to focus on the nearest layer of cells in those cases.

To quantify the regularity of dark cell and oil body patterns, we used two different spatial metrics: the proportion of idioblasts, that is, the percentage of dark cells/oil body cells that are isolated from cells of the same type; and the pattern autocorrelation through Moran’s *I* statistic (Moran, 1950), defined as:

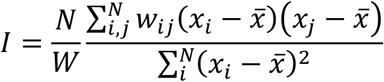

where *N* is the total number of cells in the sample, *x*_!_ are the values associated with each cell (*x*_!_ = 1 for dark/oil body cells, *x*_!_ = 0 for epidermal cells), *x̅* = ^&^ ∑^#^ *x*_!_ is the average value of *x*_!_ (equivalent to the density of dark/oil body cells), 𝑤_!"_ is the matrix of distances between cells *i* and *j*, and W = ∑^’^ 𝑤_!"_ is a normalizing factor. Typically, Moran’s *I* ranges from -1 to 1. Values close to 1 indicate positive spatial autocorrelation or clustering, while values close to -1 indicate negative spatial autocorrelation or dispersion. Values around 0 indicate randomness, without any regularity in the spatial pattern.

To test the hypothesis of increased frequency of dark cells near the edges of the thallus wings, the 55 *M. sharonae* images were narrowed down to those that included an entire thallus margin opposite the midrib. Ten images fit this criterion. For each specimen, we measured the distance of each oil body from the costa relative to the thallus edge. Because the edge is curved and the fossil thalli were not all the same size, we used distance ratios ranging from 0% (dark cells directly on the costa) to 100% (dark cells on the outer edge of the wing). It should be noted that the first row of cells adjacent to the costa are generally of a transitional shape and are very thin-walled, and sometimes a few cells were missing because the thalli tend to pull away from the costa. For these reasons, we measured distances from the first row of ’normal-shaped’ cells.

## Results

### General distribution of dark cells in M. sharonae shoots

The vast majority of *M. sharonae* specimens that were isolated from the shale possess the scattered, dark cells in both the midrib (comprised of 2-3 layers of elongate cells) and in the unistratose wings of the thallus. The dark cells are more numerous in younger tissues, such as in new branches budding off the main thallus (Fig. 2B,D) and branch apices (Fig. 2A), and at the base of more mature ventral branches (Fig. 6C). Very occasionally, specimens lack dark cells, and some have cells with varying shades of gray (Fig. 2A except for apex). Sometimes it was difficult to determine whether the few specimens that appeared to lack the cells had originally possessed them. There was variation in apparent density of dark cells and occasionally, *M. sharonae* specimens exhibited an uncharacteristically large number of dark cells, often in combination with other atypical features. For example, part of a thallus fragment might have more than 50% dark cells, while other areas might have fewer dark cells, separated by an irregular border. (Fig. 2C, E).

**Figure 2.**
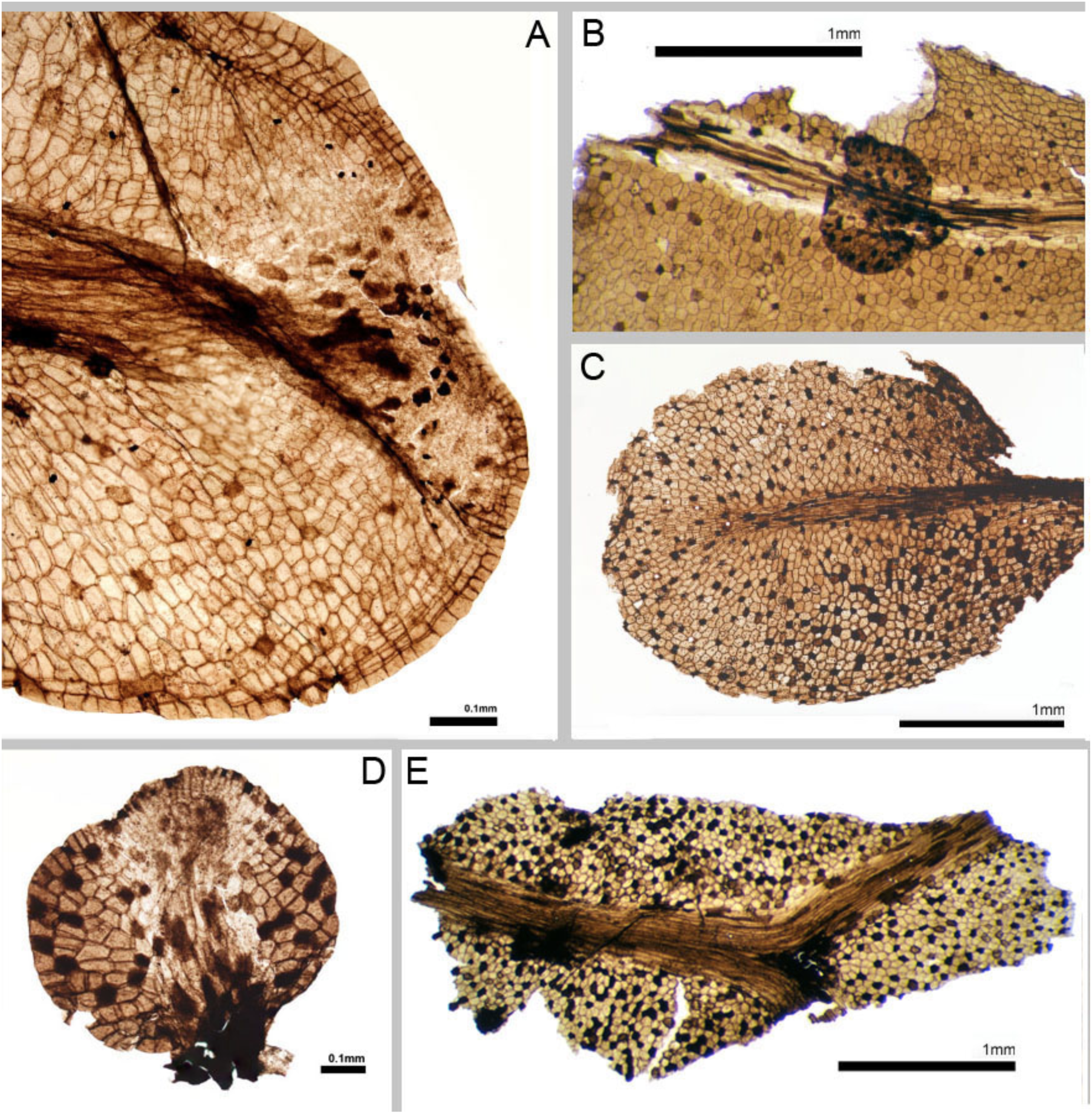
Variation in distribution of dark cells of fossil taxon *Metzgeriothallus sharonae.* A, Shoot apex with more concentrated and opaque dark cells in growing region (UC 250155.32); B, Young thallus growing from midrib has higher concentration of dark cells than the parent thallus (UC 254933.01); C, Clearly defined region in lower right part of thallus has more dark cells (UC 254936.01); D, New thallus branch (UC 254931.01); E, Fragment with many dark cells and other dark material in additional cells (UC 254918.39).

**Figure 3.**
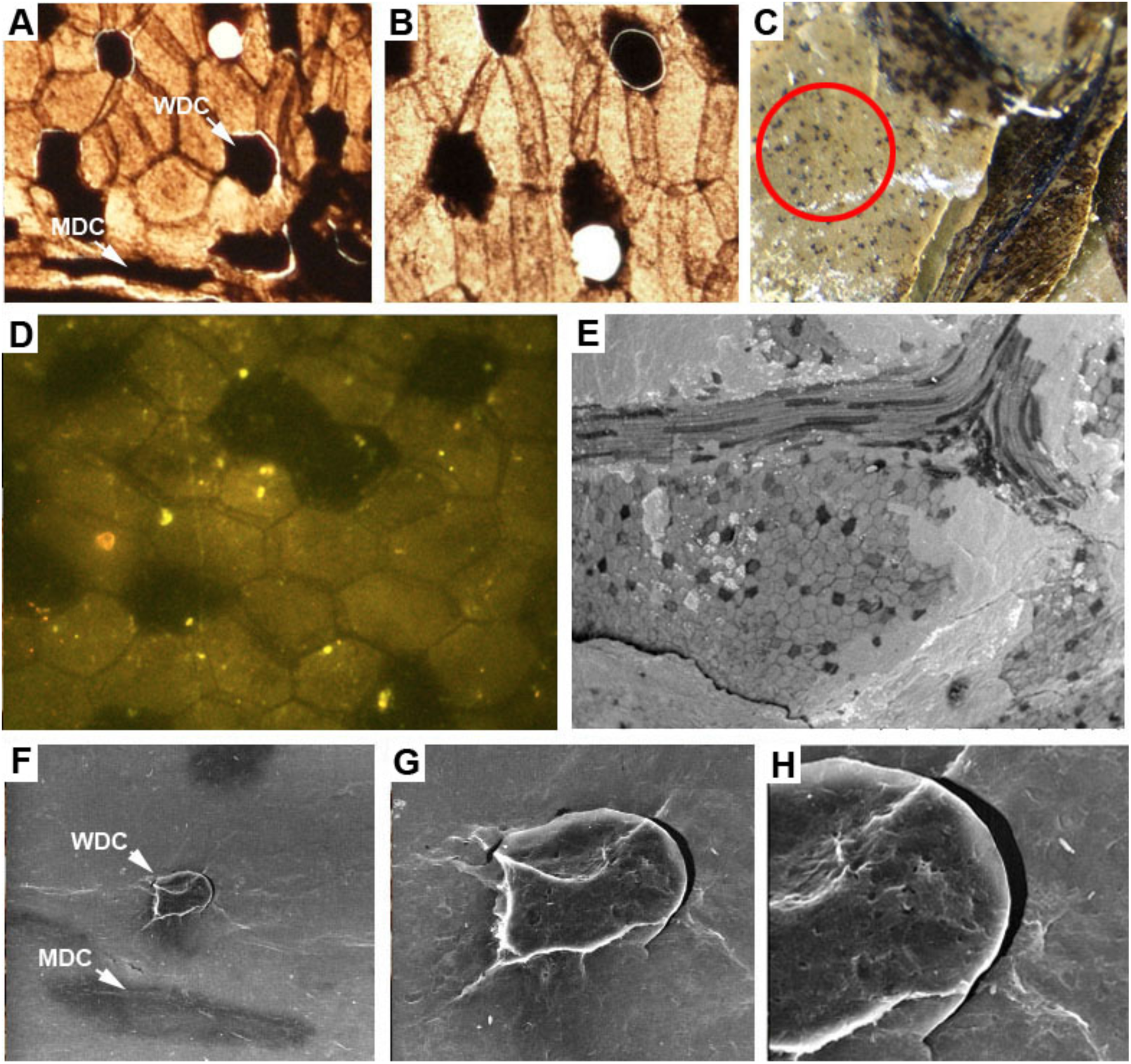
Characterization of dark cells of *Metzgeriothallus sharonae;* A, B, Circular center of dark cells partially or completely separated from fossil visible in light microscopy; C, Surface of shale matrix under polarized light showing thallus on right, on left are remains of dark cells only (circled); D, Weak, yellowish autofluorescence is confined to light cells; E, Scanning electron micrograph of uncoated surface of shale fragment near the region of a branch dichotomy; F, G & H, SEM of a dark cell in the thallus wing that is beginning to separate WDC=wing dark cell, MDC=midrib dark cell.

**Figure 4.**
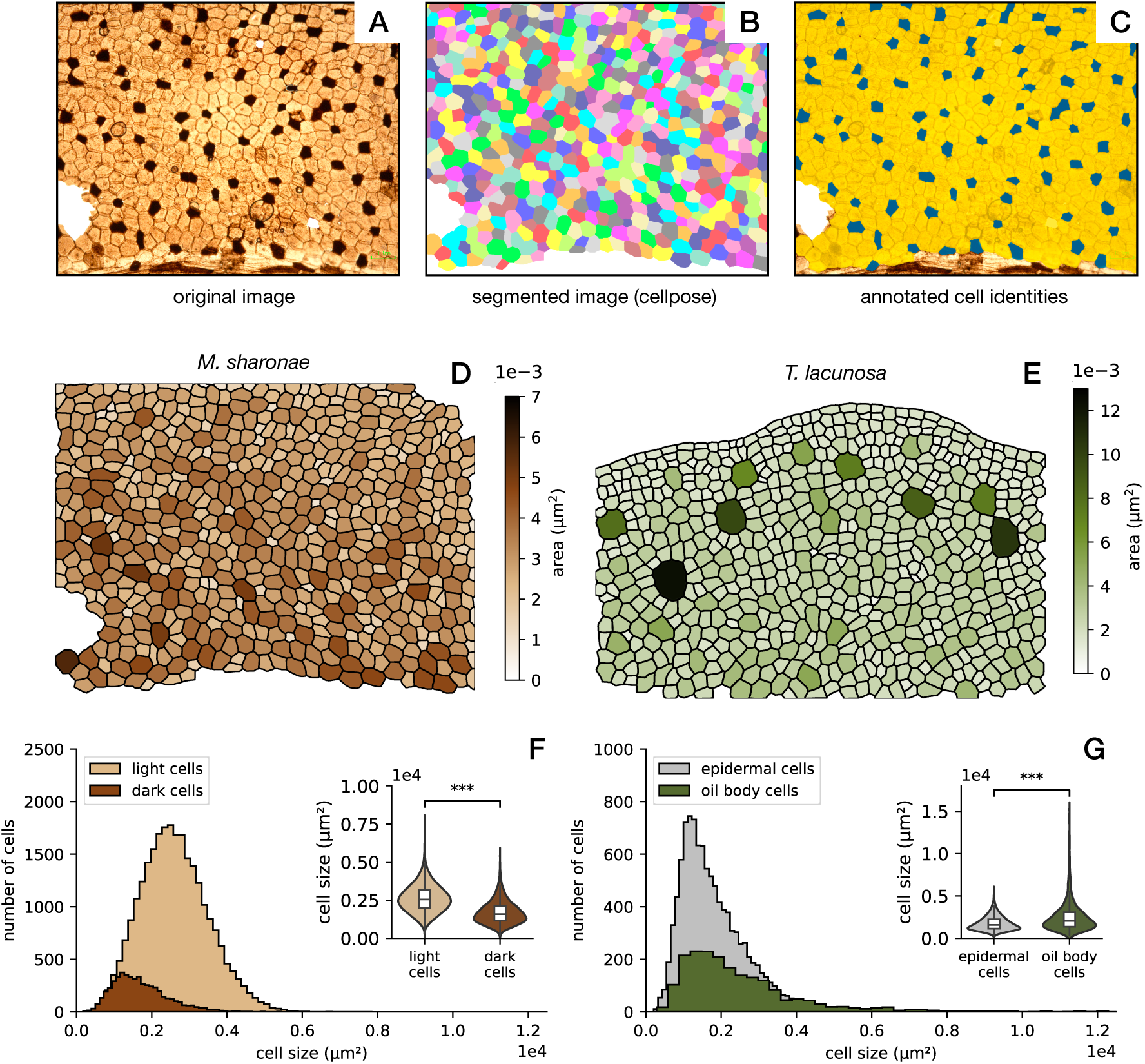
Cellular analysis of dark cells and oil body cells in *M. sharonae* and *T. lacunosa*. A-C) Segmentation pipeline for cellular analysis. For every image, cells are segmented with cellpose and subsequently labeled by type (in panel C, dark cells are blue, light cells are yellow). This pipeline allows a systematic quantification of cell sizes. D-E). Representative segmentations of *M. sharonae* (D) and *T. lacunosa* (E), coloring cells according to their area. F-G) Distribution of cell sizes in *M. sharonae* (F) and *T. lacunosa* (G). In *M. sharonae*, dark cells are on average smaller than light cells, while in *T. lacunosa*, oil body cells are on average bigger. However, in *T. lacunosa* the size distribution of oil body cells has a long tail, with some cells becoming very large. Statistical significance was computed with the Mann-Whitney U-test; ***p < 0.01.

**Figure 5.**
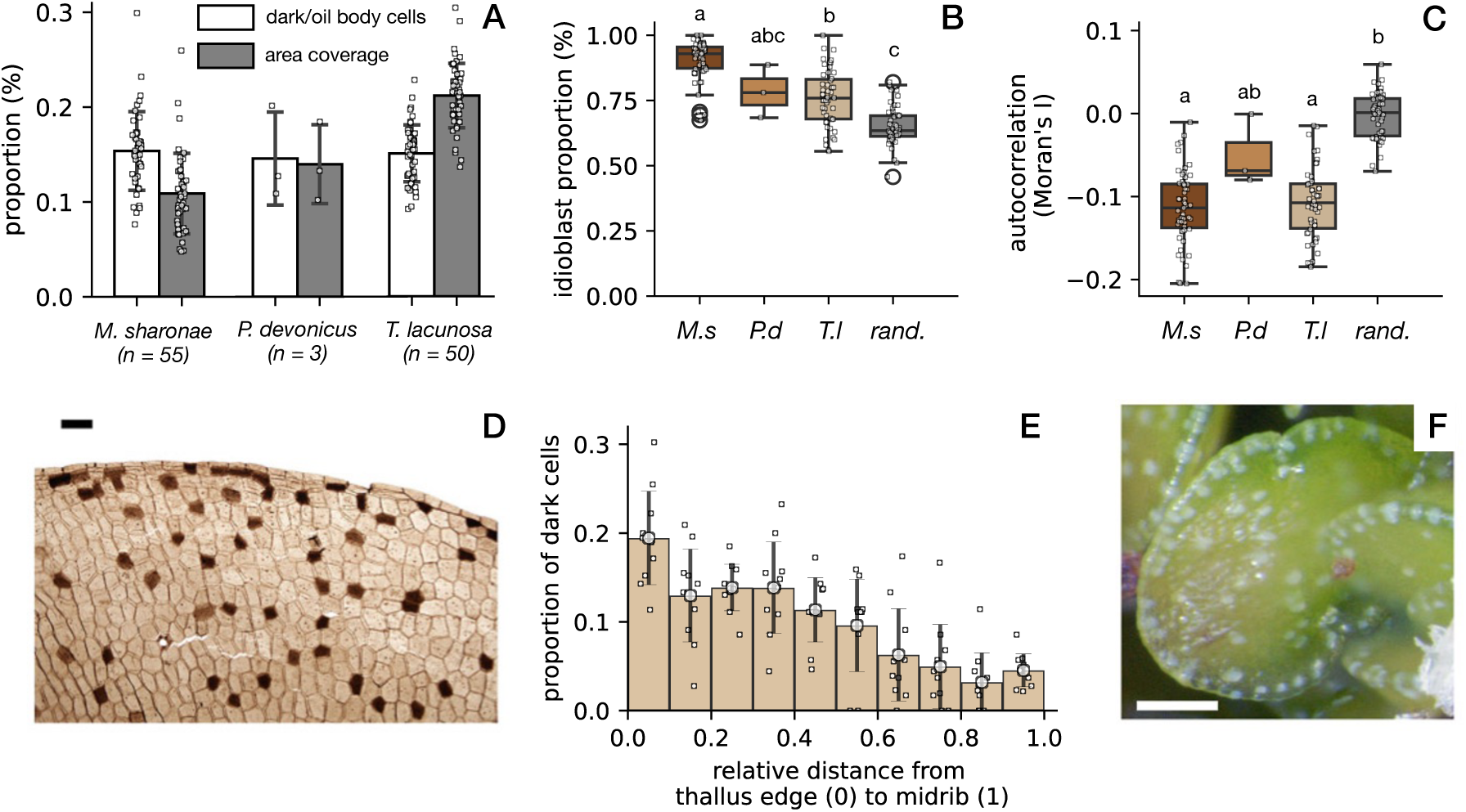
Spatial statistics of dark and oil body cells. A) Cell proportion and percentage of surface area taken up by dark cells (fossils) and oil body cells (extant taxa) or vs the total area for the four *T. lacunosa* populations and the fossil taxon *M. sharonae.* B) Idioblast proportion of fossil and extant taxa compared to what is expected if the same proportion (∼0.15) of dark/oil body cells is randomly distributed across the tissue (rand.). C) Moran’s *I* statistic of dark/oil body cell patterns, showing a significant tendency toward negative values, verifying the dispersion between dark/oil body cells. Statistical differences were tested using ANOVA followed by Tukey HSD p < 0.01; letters indicate statistically significant groups. D) Light micrograph showing a portion of unistratose wing of *Metzgeriothallus sharonae* with apparent concentration of dark cells near the thallus edge (UC 254913.03). E) Frequency distribution of the relative distance of dark cells from the thallus wings of *M. sharonae* to the midrib, obtained by combining all data points for ten specimens (812 dark cells total) and assigning the;m to ten bins. F) In the extant liverwort *Marchantia polymorpha*, oil body cells are concentrated near the edges of asexual propagules. The oil bodies of *M. polymorpha* appear white in reflected light. Scale bars 0.1mm.

**Figure 6.**
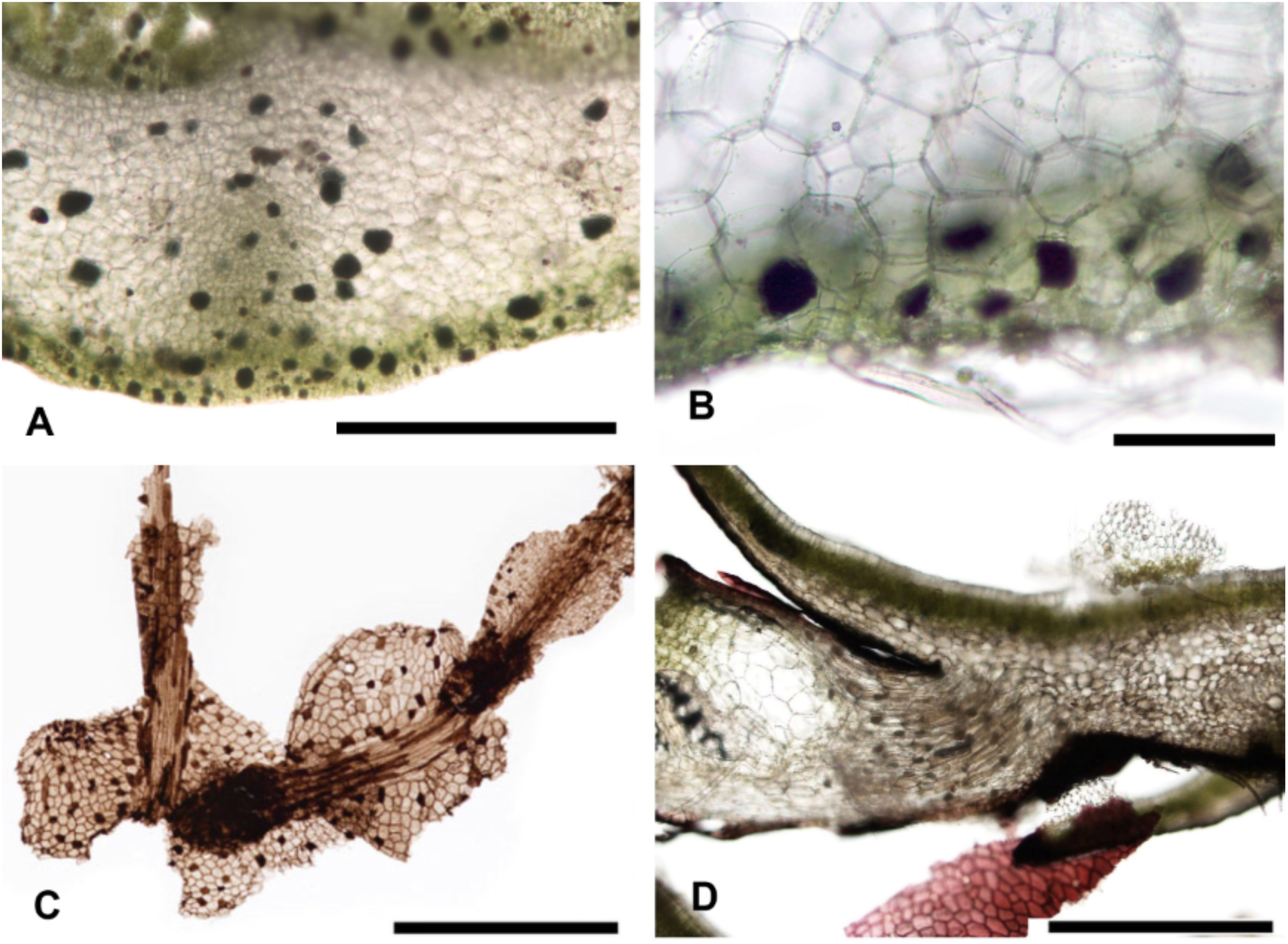
Concentration of oil body cells (extant liverworts) and dark cells (fossils) in epidermal issues and branching points. A, Cross section of *Treubia lacunosa* axis showing oil body cells concentrated near surface and around central region near concentration of fungal endophytes (UC 2091244); B, Lower surface of *Monoclea forsteri* (UC 2091243); C, Fossil taxon *Metzgeriothallus sharonae* with two ventral branches with concentration of dark cells at branch bases (UC 254913.04); D, Section of *Targionia hypophylla* showing concentration of oil body cells at base of ventral branch point (UC 2091252). Scale bars A, C, D, 1mm; B, 0.1mm

### Characterization of dark cells of M. sharonae (microscopy)

The dark cells of both costa and wing are partially to completely opaque in transmitted light. In some specimens there are round holes in the fossils that occur within the dark cells; in other cases, portions of the dark cells are partially separated but still remained attached. The holes, or dis-attached portions, form nearly perfect circles, even when they occur in the greatly elongated cells of the midrib (Fig. 3A,B). Conversely, when observing the shale surface with polarized light, black dots are sometimes visible in a similar arrangement as the dark cells of nearby fossil thalli (Fig. 3C).

### Scanning Electron Microscopy

When the uncoated fossils were imaged with SEM, the dark cells appeared consistently darker than other cells even though there was little to no variation in surface height (Fig. 3E). Figures 3F-H show increasingly close views of a portion of one of the dark cells that is breaking away from the rest of the fossil. The part that is pulling away is almost perfectly circular, suggesting that it may represent an earlier stage of the process producing the circular holes in the dark cells.

### Fluorescence Microscopy

The dark cells did not autofluoresce at all, while the light cells autofluoresced only very faintly in the yellow-orange part of the spectrum (Fig. 3D; For comparison the much more brightly autofluorescing spots in Figure 3D likely represent dust and modern contaminants of organic origin).

### Comparison of dark cells (fossils) and oil body cells (extant taxa)

In *M. sharonae*, dark cells are smaller than other cells (Fig. 4D,F). Most oil body cells in *T. lacunosa* have similar sizes than epidermal cells, but a small fraction acquire disproportionately large sizes (Fig. 4E,G) The proportion of dark cells of the two fossil taxa *M. sharonae* and *P. devonicus* and oil body cells of extant taxon *T. lacunosa* shows a high degree of conservation both within various *T. lacunosa* populations and across all taxa (Fig. 5A). All *T. lacunosa* populations showed a frequency of oil bodies within a range of 10-20%. *M. sharonae* and *P. devonicus* both had median frequencies in the same range although the sample size of *P. devonicus* was only three specimens. The area coverage of dark cells was comparable between *M. sharonae* and *P. devonicus,* but slightly higher for oil body cells in *T. lacunosa* due to their larger average size (Fig. 5A).

In most *M. sharonae* thalli, dark cells appear spatially dispersed across the tissue (Fig. 4A), with a significantly higher proportion of idioblasts than would be expected under a random distribution (Fig. 5B). Their arrangement also shows signs of spatial regularity, as reflected by negative values of Moran’s *I* statistic (Fig. 5C), indicating that dark cells tend to avoid close proximity to one another. In specimens where dark cells are more densely packed, they sometimes form linear clusters, suggesting that spatial constraints or developmental dynamics may influence local patterning. These observations point to a non-random, possibly regulated mechanism underlying dark cell placement.

### Distribution of dark cells in the unistratose ’wings’ of M. sharonae

The dark cells in *M. sharonae* are present in a much higher frequency along the edge of the fossil thalli compared to elsewhere in the thallus wings (Fig. 5D,E). The distribution of dark cells in the remainder of the thallus is relatively even, at least in terms of ’lateral’ spread of dark cells between costa and edge; the ’longitudinal’ spread along the thallus was not measured, though it appears similarly evenly distributed. There is relatively little difference in frequency in any region from 0-70% away from the midrib; however, there are 2-3 times more oil bodies in the 30% of the wings closest to the edge, and the last 1/10 of the graph shows a very large peak indicating a much higher concentration of oil bodies along the thallus edge, than in the rest of the thallus wing (Fig. 5E). The ten thalli that were used for the distribution analysis were by necessity on the small end of the thallus size distribution, as only those specimens that showed both a portion of midrib and an entire edge in the 100x magnification photographs were used in the analysis. This suggests they may have been relatively young thalli.

## Discussion

### Characterization of Metzgeriothallus dark cells

The observations with SEM, light, and stereo microscopy are consistent with a physical and chemical similarity between the fossil dark cells and oil body cells of modern liverworts. Like the cells of most land plants, liverwort cells are largely comprised of water. In contrast, the specialized oil body cells of modern liverworts are nearly filled with one large oil body. The terpenoids, bisbibenzyls and other organic compounds constituting the main components of liverwort oil bodies are carbon rich (Asakawa, 2004). The often-total opaqueness of the fossil dark cells when seen in transmitted light suggests that far more organic material has been preserved in those cells, which is consistent with the content of the carbon-rich lipids of extant liverwort oil bodies. Furthermore, the breaking away of some material from the dark cells, in a regular circular fashion (Fig. 3A,B,F-H), suggests that these areas had a denser concentration of material that over time pulled away from the surrounding material. The SEM backscatter indicates an area of greater density with a chemical composition with a low atomic number in the dark cells as compared to the light cells which also supports the hypothesis of relatively carbon-rich contents of those cells before fossilization occurred.

Both the faint autofluorescence and its yellow orange color (Fig. 3D) are consistent with the great age of the fossils (van Gijzel 1966). However, where there is a greater concentration of organic material there would be an expectation of greater fluorescence in those areas, yet we see the opposite in these images. The light, ordinary cells that were presumably mostly water have a faint, unevenly distributed fluorescence, while the dark cells do not appear to autofluoresce at all. One explanation for this could be that the material that is fluorescing is not within the cells, but from the waxy covering on the surface of the thallus. Modern land plant cuticles are formed of a combination of cutan and cutin (van Bergen *et al*., 2004). These are among the biomacromolecules that are least subject to degradation (Martin, 1999). Because the liverworts were living on or near a water source in a sub-tropical, humid climate, they may have had only a thin cuticle on the upper surface. If the specimens were oriented under the microscope with the waxy side down, the opaque material of the dark cells would have blocked the faint fluorescence completely. Autofluoresence is also indicative of the rank of coal, so that in fossils in the final stages of coalification fluorescence is completely extinguished (van Gijzel, 1966). This may also indicate that the fossils, aside from the thin cuticle, may be completely coalified.

Some *M. sharonae* specimens had no trace of the dark cells at all, possibly because these tissues had died before deposition. Oil bodies generally disintegrate after a plant is collected and begins to dry out (Schuster,1966). The specimens with unusually dense concentrations of dark cells may represent natural variation. In some species of *Treubia* today there is a great deal of variation in oil body cell frequency (Glenny *et al*., 2015).

### Spatial patterning of dark cells and oil body cells

Dark cells in *M. sharonae* are spatially distributed into regular patterns, as evidenced by the high proportion of idioblasts and the negative values of Moran’s *I* statistic (Fig. 5B,C). Regular spacing of distinct cell types is a distinctive hallmark of lateral inhibition, a mechanism in which cells that adopt a particular fate inhibit their neighbors from doing the same (Lanford *et al*., 1999; Kageyama *et al*., 2008). Such interactions typically produce "salt-and-pepper" patterns, where differentiated and undifferentiated cells are interspersed in a periodic, non-random manner (Collier *et al*., 1996, Cohen *et al*., 2023). The spatial arrangement of dark cells in *M. sharonae* closely resembles these regular patterns, suggesting that lateral inhibition may underlie their differentiation. In plants, lateral inhibition has been shown in the specification of several epidermal cell types, including leaf trichomes and root hairs in *Arabidopsis* (Schellman *et al*., 2002, Lee *et al*., 2002, Ishida *et al*., 2008), and rhizoids in *Marchantia* (Thamm *et al*., 2020, Mercadal *et al*., 2026). Our results raise the possibility that a similar mechanism governed the differentiation of oil body cells in *M. sharonae*.

Interestingly, the patterns of oil body cells in *T. lacunosa* also exhibit a high proportion of idioblasts and negative values of Moran’s *I* (Fig. 5B,C), indicating a non-random, spatially regulated distribution. A similar pattern can be observed in *Marchantia polymorpha*, where oil bodies are preferentially found in idioblasts that lack immediate neighbors with oil bodies (Fig. 5E; Romani *et al*., 2020), reinforcing the idea of spatial separation. Together, these observations further support the homology of dark cells with oil body cells, and suggest that a mechanism of patterned cell-type specification has been conserved since at least the Devonian period to regulate oil body cell differentiation. This long-term conservation points to the developmental and possibly ecological significance of positioning oil body cells in a dispersed manner within the tissue.

### Comparison with modern taxa

The distribution of specialized oil body cells in the individual plants of the Marchantiales and other groups that possess these cells has not been described in the literature beyond presence/absence in a particular taxon, as oil bodies are chiefly used taxonomically in the Jungermanniopsida. Therefore, direct observational comparisons were made with local complex thalloid taxa. The concentration of dark cells around *M. sharonae*’s branch bases (Fig. 6C) and margins of laminar plant structures (Fig. 5D) can be compared to similar observations in modern taxa such as the ventral branch bases of the modern taxon, *Targionia hypophylla* (Fig. 6D) and the gemmae (asexual propagules) of *Marchantia polymorpha* (Fig. 5F), respectively. The concentration of dark cells in younger, growing areas of *M. sharonae* such as shoot apices and new branch buds (Fig. 2A,B,D) is consistent with the hypothesis of their homology with the oil body cells; oil bodies are known to disappear in older tissues in extant liverworts, and in some taxa, such as the leafy liverwort *Lepicolea attenuata,* they are only visible in young leaves (Stewart, 1978).

Our quantitative comparisons show that the frequency of oil body distribution is highly conserved between and within the four *T. lacunosa* populations and the two Devonian fossil taxa and may imply a conserved function. Our *T. lacunosa* counts were slightly higher than those found by Glenny *et al*. (2015). This could be a result of the different methods used. Rather than count a specific number of cells, as they did, we counted all the cells in a squared-off area. Additionally, we counted into the multistratose region in order the use a similar area of tissue for all taxa, and although we were careful to count only one layer, that turned out to be difficult in practice, possibly leading to a slight overcount. Adding area coverage of specialized oil body cells to the comparison (Fig. 5A) provided an opportunity to add an example from another major lineage, the multistratose Marchantiales. It should be mentioned that *Treubia* species vary considerably in oil cell frequency (Glenny *et al*., 2015) so these results are not representative of all extant liverwort taxa with specialized oil body cells. Of the Treubiacea, *Apotreubia nana* most closely resembles the *M. sharonae* fossil liverworts in oil cell distribution, especially in the lack of size dimorphism and greater unistratose area in ’leaf’ tips (Inoue & Hattori, 1954). Unfortunately, it was not possible to obtain *A. nana* specimens for this study. The differences among three Treubiacae species are illustrated in Figure 7.

**Figure 7.**
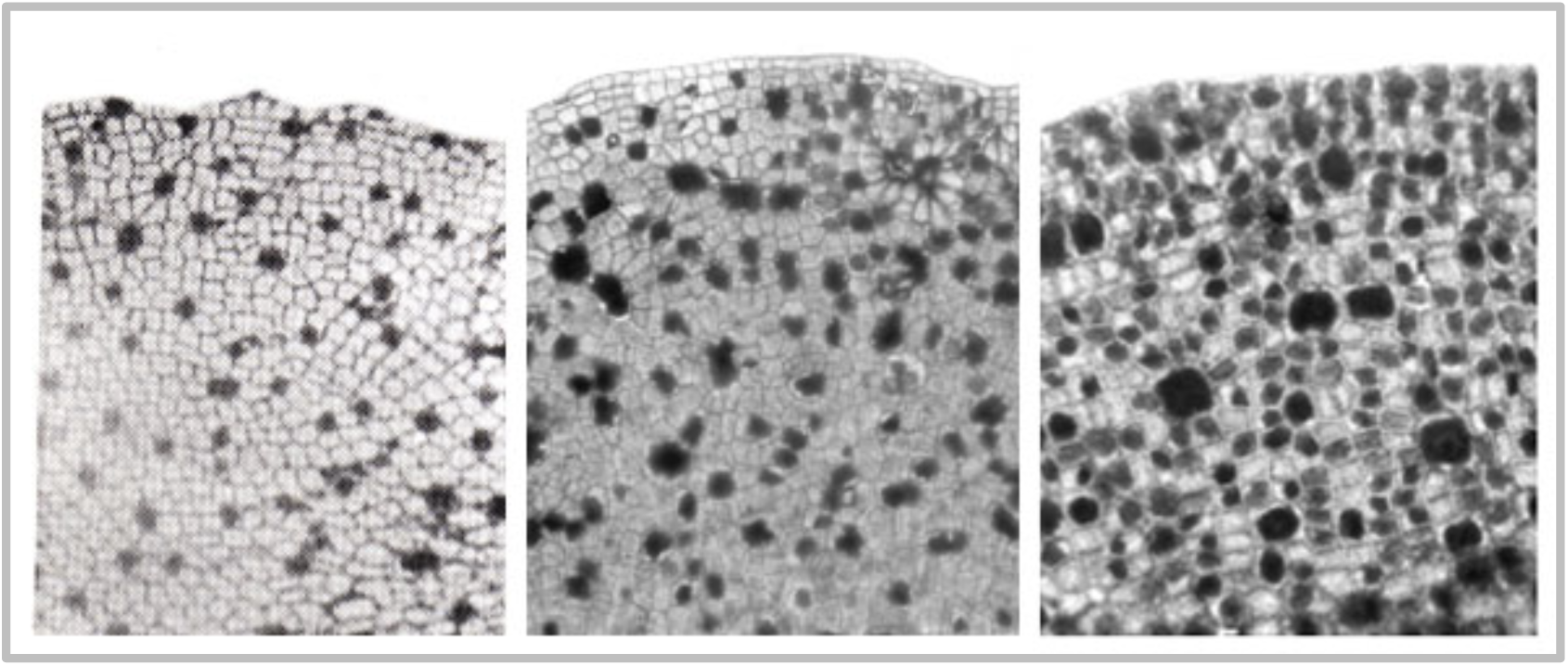
Diversity of oil body cells in ’leaves’ of extant Treubiales. Left, *Apotreubia nana* (from Inoue and Hattori 1955); Middle, *Treubia lacunosa* (UC 2036933); Right, *Treubia pygmaea* (UC 2036944).

Neither of the modern lineages in which the two Devonian fossil taxa have been systematically placed possess specialized oil body cells today (Schuster, 1966; Crandall-Stotler *et al*., 2005; Forest *et al*., 2006). Modern liverworts in the Metzgeriaceae have secondarily lost oil bodies, or on rare occasions possess numerous, minute ’oil droplets’ in each cell (Schuster, 1992) while their close relatives have oil bodies in all or most cells. In the family Pallavicinaceae, oil bodies are found in most cells of the plant, with each cell usually containing several oil bodies rather than one large oil body (Schuster 1966). This puts specialized oil body cells in all three main lineages: in extant taxa in the Haplomitriopsida and Marchantiopsida and fossil taxa of the Jungermanniopsida.

### Anti-herbivory hypothesis

The results of the analysis of the distribution of dark cells of *Metzgeriothallus* between midrib and thallus edge (Fig. 5D,E) confirm the observation of the non-random, even distribution of the dark cells of the fossil, as well as their increased frequency near the edge of the vegetative thalli (Labandeira *et al*., 2013). These results support the suggestion by Labandeira *et al*. (2013) of a possible early oil body anti-herbivory function.

The observations of increased frequency of the fossil dark cells in growing areas and young thallus branches also supports the anti-herbivory hypothesis of oil body function, as these young tissues are more vulnerable to predation. The anti-herbivory hypothesis has experimental support in the modern liverworts (He *et al*., 2013; Romani *et al.,* 2020; Gutsche *et al*. 2024) and is also supported by results that show that terpenoids similar to those found in liverwort oil bodies have anti-herbivore effects in vascular plants (Lange 2015, Tanaka *et al*. 2016). It is worth noting that in the multistratose regions of *Treubia lacunosa* and *Monoclea forsteri* most oil body cells are concentrated in the cell layers near the upper and lower epidermis (Fig. 6A,B) which may indicate a similar protective function.

The morphological, chemical, and quantitative results support the hypothesis that the dark, scattered cells of *Metzgeriothallus sharonae* and *Pallavicinites devonicus,* and other Paleozoic liverworts that possess these cells, are homologous to the specialized oil body cells of extant liverworts. These results provide an important new character that can be used in assigning fossils and in elucidating oil body evolution and function.

## Acknowledgements

Funding for this research was provided by the University of California, Berkeley’s Department of Integrative Biology, the Erwin Resetko Family scholarship fund and the Evolving Earth Foundation. We thank NMNH for loan of the original material collected by Fran Hueber, and ST thanks David Glenny and Bill and Nancy Malcolm for hospitality and support in the field in New Zealand. Taehee Han and Junnan Chen provided laboratory assistance. JM thanks Pau Formosa-Jordan for hosting him at the MPIPZ, where this work took place. JM acknowledges financial support from the Alexander von Humboldt-Stiftung (ESP 1236193 HFST-P) and the European Union’s Horizon 2023 research and innovation programme under the Marie Skłodowska-Curie grant agreement No 101153033.

## Author contributions

ST provided research design, fossil collection and processing, field work and plant collection, and microscopy and photography. ST wrote the manuscript with the help of JM. JM performed image segmentation and conducted the statistical and data analyses.

## Data Availability

The code required for data analyses presented in this paper is available upon request.

